# Research disturbance negatively impacts incubation behaviour of female Great tits

**DOI:** 10.1101/2024.03.28.587136

**Authors:** Léanne Clemencin, Emilio Barba, David Diez-Méndez

## Abstract

Human-induced disturbance is perceived by avian species as a predation risk. However, the anti-predatory behaviour triggered by these non-lethal events can have negative impacts on reproduction and offspring survival. Research on breeding birds often involves visits to their nests and is likely to disrupt parental behaviour, but nest visits that do not involve direct handling of females have been overlooked as important disturbance events. This study focuses on the impacts of short visits to the nest of incubating Great tit *(Parus major*) females. We investigated how long they stay away from the nest (off-bout) after a disturbance, their possible compensatory behaviour once they resume incubation (on-bout), and the effects on daily incubation rhythms. We used three years of data from two breeding populations to assess the consequences of disturbances in two scenarios: when the female is present in the nest and flushed, and when the female is absent. We found that after a disturbance, the immediate off-bout was longer when the female was either present or absent, with the magnitude of the disturbance being greater when females were flushed. Females did not compensate with longer on-bouts afterwards, i.e. the research disturbance altered daily incubation behaviour by reducing the total time spent on the nest in relation to the number of daily disturbance events. Females that alter their behaviour in response to perceived predation risk would perform longer incubation periods, resulting in lower hatching rates. These effects of research on female behaviour should be considered when planning field experiments.

## Introduction

Human-induced disturbance events to avian nests are perceived similarly to predation risk (“risk-disturbance hypothesis”, Frid and Dill 2002), although they are usually non-lethal by nature (Cresswell 2008). In response, animals may adjust their behaviour (Beale and Monaghan 2004; Perona et al. 2019), physiology (Eggers et al. 2006; Coslovsky and Richner 2011) or life-history traits (Hutfluss and Dingemanse 2019) to the magnitude of perceived predation risk (Lima and Bednekoff 1999). Anti-predatory behavioural responses can have a negative impact on reproduction and offspring survival, even if they contribute to a breeding individual not being predated. During egg-laying and incubation, parents that flee the nest in response to perceived risk facilitate nest predation (Stien and Ims 2016). Even when predation does not occur, the absence of parents alters incubation rhythms (Verhulst et al. 2001; Baudains and Lloyd 2007) and increase temperature fluctuations in the clutch (DuRant et al. 2013), which could have a long-term negative impact on offspring survival (Berntsen and Bech 2016). After egg hatching, nest disturbance also affects parental care by reducing their feeding activity (Verhulst et al. 2001) and impairing offspring growth (Remacha et al. 2016). Such impacts resulting from perceived predation risk are often associated with human recreation (Watson et al. 2014; Perona et al. 2019), although research activities during the breeding season may have analogous effects (Stien and Ims 2016).

Nest disturbance is inevitably associated with studies of avian reproduction (Lenington 1979). It can be assumed that research activities are more disruptive than recreational activities, as they usually involve handling individuals, visiting nests or manipulating eggs, even though only temporarily. Attracting predators to nest sites has traditionally been the main impact that researchers have been concerned about. Although an early review showed no clear patterns overall (Götmark 1992), there was high interspecific variation. Later studies not only found no measurable negative effects (Schaub et al. 1992; Lloyd et al. 2000; Weidinger 2008; Hu et al. 2020) but even positive effects of research activities (Ibáñez-Álamo and Soler 2010), as the presence of humans near nests may deter mammal predators rather than attracting them (reviewed in Ibáñez-Álamo et al. 2012). However, as with recreational activities, nest predation is only one possible effect, and other consequences of research activities on breeding individuals may also be at play, such as altered incubation patterns and even nest abandonment (Dubiec 2011).

Visiting a nest during the incubation period is a research activity which impact has rarely been assessed (see also Croston et al. 2021), probably because it is considered minimal. The incubation period is of particular interest because females, in gyneparental species, spend up to 80% of their active time at the nest incubating (on-bouts), leaving only briefly for foraging or other self-care activities (off-bouts) (e.g., Ardia et al. 2010; Johnson et al. 2013; Ospina et al. 2018; Diez-Méndez et al. 2021a). Given the extent of nest attendance, there is a high probability of encountering the incubating female on a regular visit to the nest. As females are largely constraint to maintain an adequate clutch temperature for embryonic development (Nord and Nilsson 2011; DuRant et al. 2013), at a high energy cost (Nord and Williams 2015), incubating females could be expected to abandon their nests when predation risk is considered high. This is particularly true in the first half of the incubating period when, according to life-history theory, deserting the nest may provide higher fitness benefits in the future than further investment in the current reproductive event (Clutton-Brock 1991; Stearns 1992). For this reason, assessments of research activity in this period have only been conducted when visiting a nest implies high desertion rates, like after handling incubating females (up to 41% of desertions, Kania 1989; Kilgas et al. 2007; Dubiec 2011) or after experimental drastic reductions in clutch size (e.g., simulating partial predation, Verboven and Tinbergen 2002; Johnston 2011; Zieliński 2020). In contrast, research on the effects of research visits that do not imply handling, is lacking.

Visiting a nest and flushing the female to determine clutch size, a common practise, may also affect incubation behaviour, but to a lesser extent than the above-mentioned disturbances. Simply visiting a nest appears to be perceived by females as a predation risk and triggers anti-predator behaviours such as hissing (Krams et al. 2014; Zub et al. 2017; Dutour et al. 2020; van Iersel et al. 2023). Previous studies using stuffed predators, mounted outside the nest-box to simulate predation risk, resulted in an overall decrease in nest attendance: females reduced their activity by returning later after being exposed to the predator (longer off-bouts) and limiting the time they spent in the nest (shorter on-bouts) (Zanette et al. 2011; Ibáñez-Álamo and Soler 2012). The effects of a disturbance event can last up to 48 hours afterwards (i.e., recovery time, Tkaczyk et al. 2023). Repeated prolonged absence of the incubating female may have dramatic consequences for embryonic development as temperature fluctuations increase (Olson et al. 2006) and the incubation period becomes longer, resulting in reduced egg hatchability (Nord and Nilsson 2011; Macdonald et al. 2013; Diez-Méndez et al. 2020) and post-fledging survival rates (Hepp et al. 2006; Berntsen and Bech 2016; Ospina et al. 2018, reviewed in DuRant et al. 2013). However we cannot ignore other studies that also used stuffed predators and found that the observed changes in the incubation rhythm, i.e., lengthening both off- and on-bouts proportionally, did not reduce nest attentiveness (Conway and Martin 2000; Ghalambor and Martin 2002; Kovařík and Pavel 2011; Basso and Richner 2015a; Tkaczyk et al. 2023, but see Basso and Richner 2015b).

Here we investigated the effects of research disturbance during the incubation period in two Great tit (*Parus major*) breeding populations. We assessed the effects of disturbance and potential recovery time in two common scenarios when conducting nest visits: 1) the incubating female is present in the nest (on-bout) and is consequently flushed, and 2) the female is absent (off-bout). Female flushing has been dismissed as an important disturbance in this species because, to our knowledge, its effects have not been considered so far, despite Great tits being one of the most common model species in the wild. We hypothesized that visiting a nest-box would have a negative effect, albeit limited, on incubation behaviour in both scenarios, i.e., either when the female is present and flushed or when she is absent. A limited negative effect means that we would expect a short recovery time, with females lengthening just the immediate off-bout after a disturbance event, to be followed by a longer on-bout to compensate. In addition, some visits to the nest lasted longer than others, depending on the ongoing experiments (e.g., weighing and measuring individual eggs), and we anticipated to find stronger responses in nests that we had disturbed for longer. We expected that these changes in the incubation rhythm would not cause overall changes in daily incubation behaviour (i.e., nest attentiveness) even if we visited a nest twice a day. We hypothesised that the effect of research disturbance in the absence of a female would still elicit a noticeable behavioural response because foraging territories are small (Naef-Daenzer and Keller 1999) and females can still detect intruders. We also expected that flushing a female triggers a stronger response, i.e., longer off-bouts and proportionally longer on-bouts, than when females are absent as it may be perceived as a riskier encounter.

## Material and Methods

### Study area and species

Great tits are small (12.5 – 15 cm) secondary hole-nesting passerines that naturally breed in cavities but also readily accept nest-boxes (Perrins 1979). They lay one egg per day and usually begin incubating when the clutch is complete, although there is great variation depending on external conditions. The incubation period usually lasts 12-13 days (Diez-Méndez et al. 2021b). They are assisted gyneparental incubators (Nord and Williams 2015), which means that only the females incubate and the male may assist by bringing food to the nest (Bambini et al. 2019). Nevertheless, the female regularly leaves the nest to forage and perform other self-care activities (intermittent incubation).

Breeding data was collected during three consecutive breeding seasons (2015-2017) in two Mediterranean nest-box populations from eastern Spain: Sagunto and Pina. Data collection was part of ongoing long-term projects running since 1987 and 2007 respectively (e.g., Álvarez et al. 2013). Sagunto (39.70°N, 0.25°W) contains 500 nest-boxes. Sagunto is a highly anthropized habitat part of an extensive monoculture of orange trees (*Citrus aurantium*) at sea level (30 m a.s.l.). Pina (40.02°N, 0.63°W) contains 200 nest-boxes and is part of a mountain range (1200 m a.s.l.) which contains a mixed forest dominated by maritime pines (*Pinus pinaster*) with scattered stands of Portuguese oaks (*Quercus faginea*).

### Field procedures

We monitored the nest-boxes looking for first clutches of breeding Great tits in both populations on a weekly basis, and increased the frequency to daily when a nest was fully built to determine when the first egg was laid in each nest. In brief, after the first egg was laid, we attached a probe (30-gauge wire, accuracy ± 1.0°C; Onset Computer Corporation, USA) to the centre of the nest cup, which was connected through the bottom of the nest to a thermocouple data logger (HOBO UX100-014M Single Channel, Onset Computer Corporation, USA) placed between the nest and the bottom of the nest-box (see Diez-Méndez et al. 2021b). We recorded the egg temperature every 10s until the first egg hatched in order to infer female incubation behaviour from the temperature fluctuations in the egg. The probe was inserted into a plastic egg (Factory Direct Craft Supply, USA), filled with a lubricant (Clear Glide, Ideal industries, USA) that was similar in shape and colour to the Great tit eggs but slightly smaller (15.9 × 11.5 mm versus 18.1 × 13.3 mm; average egg size from Sagunto; Encabo et al. 2001). In addition, we attached a Thermochron iButton data logger (accuracy ± 0.5°C, model DS1922L-F5, Maxim Integrated) to the upper inner part of each nest-box wall to record the local ambient temperature every 520 s.

Each nest-box was visited a minimum of three days during the incubation period: 1) when we found an incubating female or warm eggs during a regular morning check for the first time, 2) about five days into the incubation period, when we visited the nest to check the probe and clutch size, and 3) eleven days into the incubation period, when we began monitoring egg hatching. The latter implied two visits per day, in the early morning and late afternoon. The total number of visits to a nest depended on how long the eggs took to hatch and the how the requirements of an ongoing experiment were. Each time we visited a nest, we noted the time and whether the female was present or absent. If the female was present, we always flushed her out to check the contents of the nest. The standard visits to the nest-boxes lasted less than a minute (i.e., a short disturbance), but depending on the year and ongoing experiments (e.g., measuring and weighing individual eggs), these visits lasted longer (i.e., 1-5 min, long disturbance events).

### Incubation behaviour and disturbance measurements

Incubation behaviour of Great tit females was assessed using recorded nest-cup temperatures in each nest during the incubation period, We defined the onset of incubation as the first time nest attentiveness exceeded 50% of the active day (incubation day 1), and the incubation period lasted to the day before the first egg hatched (following Cresswell and McCleery 2003; Simmonds et al. 2017; Diez-Méndez et al. 2021b). Nest attentiveness was defined as the daily sum of minutes a female spent incubating in the nest compared to the duration of the active day. A female’s active day duration was calculated as the sum of minutes between the first off-bout in the early morning and the last on-bout in the evening to overnight.

To identify off- and on-bouts, we used the software *Rhythm* (Cooper and Mills 2005) and followed the criteria of setting off-bouts to last at least 2 min, a minimum temperature decrease of 2.0°C and a minimum initial cooling slope of 0.2°C. We then visualized the Rhythm output in Raven (Cornell Laboratory of Ornithology) to check the timing and duration of the bouts. We edited the start or end of bouts when the algorithm did not cover the entire duration, discarded temperature drops misidentified as off-bouts, and included clear off-bouts that did not meet the minimum duration criteria. To minimize observer bias, blinded methods were used when temperature data were analyzed to identify off- and on-bouts. Only off-bouts with a duration of less than 120 minutes were taken into account, as longer off-bouts are rare and considered outliers (∼1.5% of total off-bouts, Diez-Méndez et al. 2021a).

### Statistical analyses

We performed statistical analyses with the software R 4.3.1 (R Core Team 2023). Six linear (LMMs) and one generalised linear mixed-effects model (GLMMs) were created using the package *lme4* (Bates et al. 2015). We built a LMM to assess the off-bout duration when a female was absent during a nest-box visit (disturbance) using the off-bout temporal sequence (before, during and after disturbance) as a fixed categorical factor together with day of incubation, ambient temperature (quadratic term, poly-function), time of the day (quadratic term, poly-function), calendar date, population and year, including nest-box as a random effect. A second LMM was built to assess the off-bout duration when a female was present in the nest during a visit, using off-bout categorization (before, immediately after the disturbance, and the next one) as a fixed factor. The third LMM looked at the duration of the on-bouts when a female was absent, using a fixed factor categorizing on-bouts before (one) and after the disturbance (two on-bouts). The fourth LMM dealt with the duration of the on-bouts when the incubating female was present during a visit to the nest, using a fixed effect to categorize on-bouts before (one) and after the disturbance (two on-bouts). We did not consider the on-bout that abruptly finished with the disturbance because its duration was shortened by the visit itself. Bout duration was measured in minutes and log transformed in every model. Two more LMMs were built to determine whether the duration of the disturbance influenced the subsequent off-bout duration. Off-bout duration when a female was present or absent was used as the dependent variable, and disturbance duration was an independent factor with two levels: short and long. The rest of the fixed and random factors were similar to the previous LMMs.

Our last model (GLMM, binomial family) dealt with daily nest attentiveness and added disturbance duration as a fixed factor. The number of disturbance events per day ranged from 0 to 2. For each day with one or two disturbance events in a given nest, we included an additional day, during the incubation period, when we did not visit that nest. In addition, day of incubation, mean daily ambient temperature (quadratic, poly-function), calendar date, population and year were included as fixed factors, and nest-box as a random factor in the model. All numerical variables were centred. Normality, homogeneity of variance and collinearity were visually checked for the LMMs models using the *performance* package (Lüdecke et al. 2021). The resulting figures were created with *ggplot2* (Wickham 2016) after calculating the estimates ± 95%CI with the package *Effects* (Fox and Hong 2009).

## Results

A total of 46 nests, 29 from Sagunto and 17 from Pina, were suitable for assessing the effects of research visits on female incubation behaviour. We recorded 273 disturbances and analysed the on- and off-bout duration before, during and after an event, totalling 1,638 bouts (Suppl. Mat. Table S1). Our data showed the general pattern of off-bouts being three times shorter than on-bouts before any disturbance altered them (median (range): ∼10 (1.8 – 116) vs. ∼30 (4.2 – 139) min, Supp. Mat. Table S1).

When the female was absent from the nest during a nest-box visit, the ongoing off-bout lasted longer than both the previous (estimate ± SE = −0.27 ± 0.098, t = −2.80, *p* = 0.006) and following off-bouts (estimate ± SE = −0.42 ± 0.096, t = −4.40, *p* < 0.001) (Table 1, Fig. 1a). We also found that off-bout duration of absent females was negatively affected by the incubation day (estimate ± SE = −0.21 ± 0.059, t = −3.58, *p* < 0.001) (Table 1). However, we found no detectable effect on the first on-bout duration after a disturbance event compared to the previous and the following on-bouts (Table S2, Fig. 1c). The duration of a disturbance had not measurable effects on the ongoing off-bout duration (Table S3), but overall the ongoing off-bout tended to be shorter in the Sagunto population (an anthropized orange plantation) than in Pina (a mixed forest) (estimate ± SE = −0.58 ± 0.303, t = −1.93, *p* = 0.060) (Table S3).

**Fig. 1.**
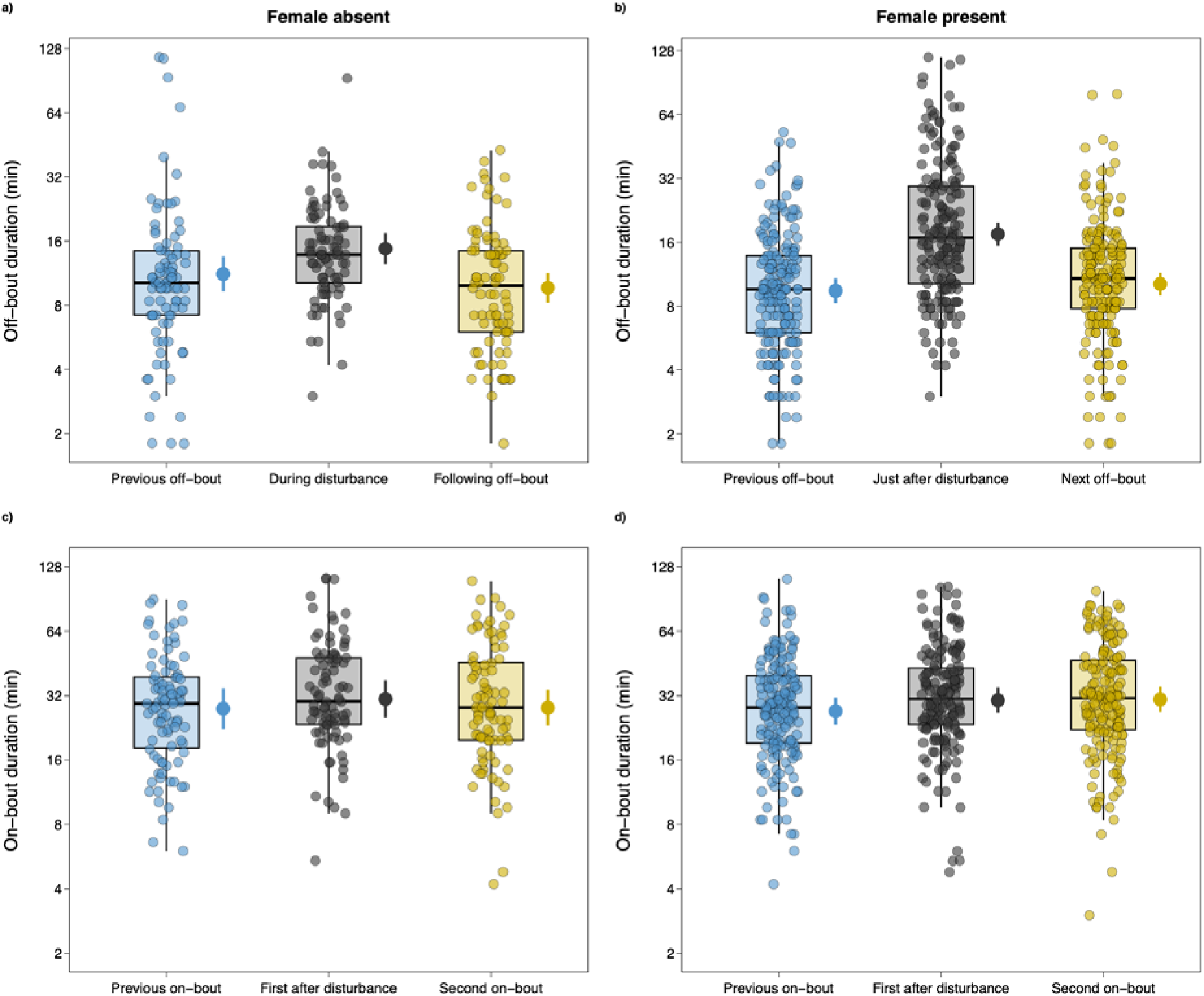
Comparison of the duration (log scale) of the ongoing/first bout after a disturbance event and the bout before the event and the next one. The incubating female may be present (on-bout) or absent (off-bout) during such events. Box-plots represent the median and 50% of the values, and upper and lower whiskers represent values > 75th and < 25th percentile. Semi-transparent circles represent the bout duration raw data. Full circles and associated bars show the model estimates and 95% confidence intervals

**Table 1.** Estimates of linear mixed-effect models analysing the effect of a disturbance event on the ongoing off-bout (female absent in the nest-box) compared to the previous and next off-bouts, or the immediate off-bout after a disturbance (female present and flushed out of the nest-box) compared to the previous and the next off-bouts. Incubation day, ambient temperature, time of the day, population and year were also considered. Nest-box is a random effect. Significant effects P < 0.05 are highlighted in bold

|  | Estimate | SE | *t* | *P* |
| --- | --- | --- | --- | --- |
| Off-bout (female absent) |  |  |  |  |
| $(R^2_{\text{fixed factors}} = 0.18, R^2_{\text{model}} = 0.23)$ | | | | |
| Intercept | 2.74 | 0.200 | 12.46 |  |
| **Previous** | **-0.27** | **0.098** | **-2.80** | **0.006** |
| **Next** | **-0.42** | **0.096** | **-4.40** | **<0.001** |
| **Incubation day** | **-0.21** | **0.059** | **-3.58** | **<0.001** |
| Temperature | 0.87 | 0.806 | 1.07 | 0.284 |
| Temperature<sup>2</sup> | -0.49 | 0.736 | -0.67 | 0.503 |
| Time | 0.27 | 0.803 | 0.33 | 0.739 |
| Time<sup>2</sup> | -0.71 | 0.857 | -0.83 | 0.408 |
| Date | 0.01 | 0.130 | 0.05 | 0.963 |
| Population: Sagunto | -0.24 | 0.244 | -0.99 | 0.330 |
| Year 2016 | 0.18 | 0.177 | 1.01 | 0.319 |
| Year 2017 | 0.03 | 0.171 | 0.19 | 0.851 |
| Off-bout (female present) |  |  |  |  |
| $(R^2_{\text{fixed factors}} = 0.20, R^2_{\text{model}} = 0.22)$ | | | | |
| Intercept | 2.75 | 0.207 | 13.32 |  |
| **Previous** | **-0.61** | **0.069** | **-8.84** | **<0.001** |
| **Next** | **-0.54** | **0.069** | **-7.84** | **<0.001** |
| **Incubation day** | **-0.11** | **0.039** | **-2.91** | **0.004** |
| **Temperature** | **2.72** | **0.856** | **3.18** | **0.002** |
| Temperature<sup>2</sup> | 0.52 | 0.718 | 0.73 | 0.466 |
| Time | 0.30 | 0.802 | 0.37 | 0.709 |
| Time<sup>2</sup> | 0.13 | 0.807 | 0.16 | 0.872 |
| Date | 0.08 | 0.084 | 1.00 | 0.323 |
| Population: Sagunto | -0.03 | 0.178 | -1.17 | 0.867 |
| Year 2016 | 0.24 | 0.152 | 1.59 | 0.117 |
| Year 2017 | 0.12 | 0.149 | 0.83 | 0.409 |

When the female was present in the nest during a nest-box visit, the immediate off-bout also lasted longer than the previous and following off-bouts (Table 1, Fig. 1b), and the two on-bouts following a disturbance event tended to be longer than before (estimate ± SE = −0.12 ± 0. 061, t = −1.92, *p* = 0.055), but the effect was not clear (Table S2, Fig. 1d). Finding a female in the nest, and flushing her out, led to a subsequent longer off-bout compared to off-bouts from females that were out of the nest. While in the latter females extended their off-bout by ∼35% on average, the former performed ∼75% longer off-bouts (Table S1). As incubation progressed, present females also shortened their off-bouts (estimate ± SE = −0.11 ± 0. 039, t = −2.91, *p* = 0.004), and lengthened their off-bouts at increasing ambient temperatures (estimate ± SE = 2.72 ± 0. 856, t = 3.18, *p* = 0.002) (Table 1). The duration of the disturbance also affected for how long a female stayed off the nest after a visit: a long disturbance caused a longer off-bout (estimate ± SE = 0.37 ± 0. 138, t = 2.69, *p* = 0.017) (Table 2).

**Table 2.**
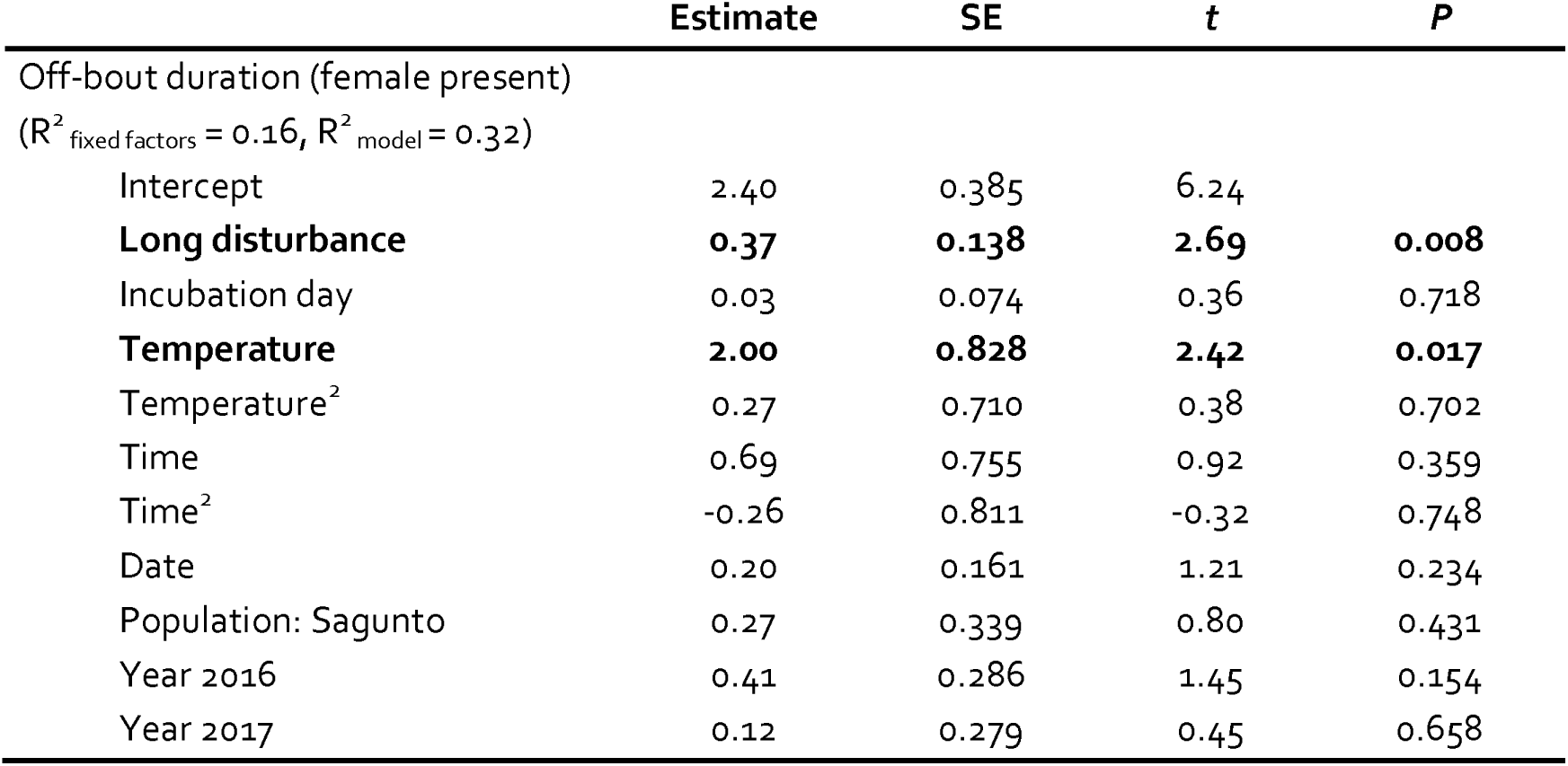
Estimates of a linear mixed-effect model analysing the effect of the duration of a disturbance event (short vs. long disturbance) on the duration of the subsequent off-bout when a female was present and flushed out of the nest-box. Incubation day, ambient temperature, time of the day, population and year were also considered. Nest-box is a random effect. Significant effects P < 0.05 are highlighted in bold

|  | Estimate | SE | *t* | *P* |
| --- | --- | --- | --- | --- |
| Off-bout duration (female present) |  |  |  |  |
| (R<sup>2</sup><sub>fixed factors</sub> = 0.16, R<sup>2</sup><sub>model</sub> = 0.32) |  |  |  |  |
| Intercept | 2.40 | 0.385 | 6.24 |  |
| **Long disturbance** | **0.37** | **0.138** | **2.69** | **0.008** |
| Incubation day | 0.03 | 0.074 | 0.36 | 0.718 |
| **Temperature** | **2.00** | **0.828** | **2.42** | **0.017** |
| Temperature<sup>2</sup> | 0.27 | 0.710 | 0.38 | 0.702 |
| Time | 0.69 | 0.755 | 0.92 | 0.359 |
| Time<sup>2</sup> | -0.26 | 0.811 | -0.32 | 0.748 |
| Date | 0.20 | 0.161 | 1.21 | 0.234 |
| Population: Sagunto | 0.27 | 0.339 | 0.80 | 0.431 |
| Year 2016 | 0.41 | 0.286 | 1.45 | 0.154 |
| Year 2017 | 0.12 | 0.279 | 0.45 | 0.658 |

Disturbances in the nest were reflected in daily nest attentiveness. Females spent less time incubating when they underwent two disturbance events in a single day (estimate = 67.4, 95% CI = 64.1 – 70.6) than when there was only one (estimate = 71.3, 95% CI = 68.7 – 73.7), and less time when there was only one daily event than when there was no disturbance at all (estimate = 76.1, 95% CI = 74.0 – 78.1) (Table 3. Fig. 2). We also found that both incubation day and daily ambient temperature increased (estimate ± SE = 0.19 ± 0. 047, t = 3.97, *p* < 0.001) and decreased (estimate ± SE = −0.91 ± 0. 340, t = −2.68, *p* = 0.007) female daily incubation effort (i.e., nest attentiveness), respectively (Table 3).

**Fig. 2.**
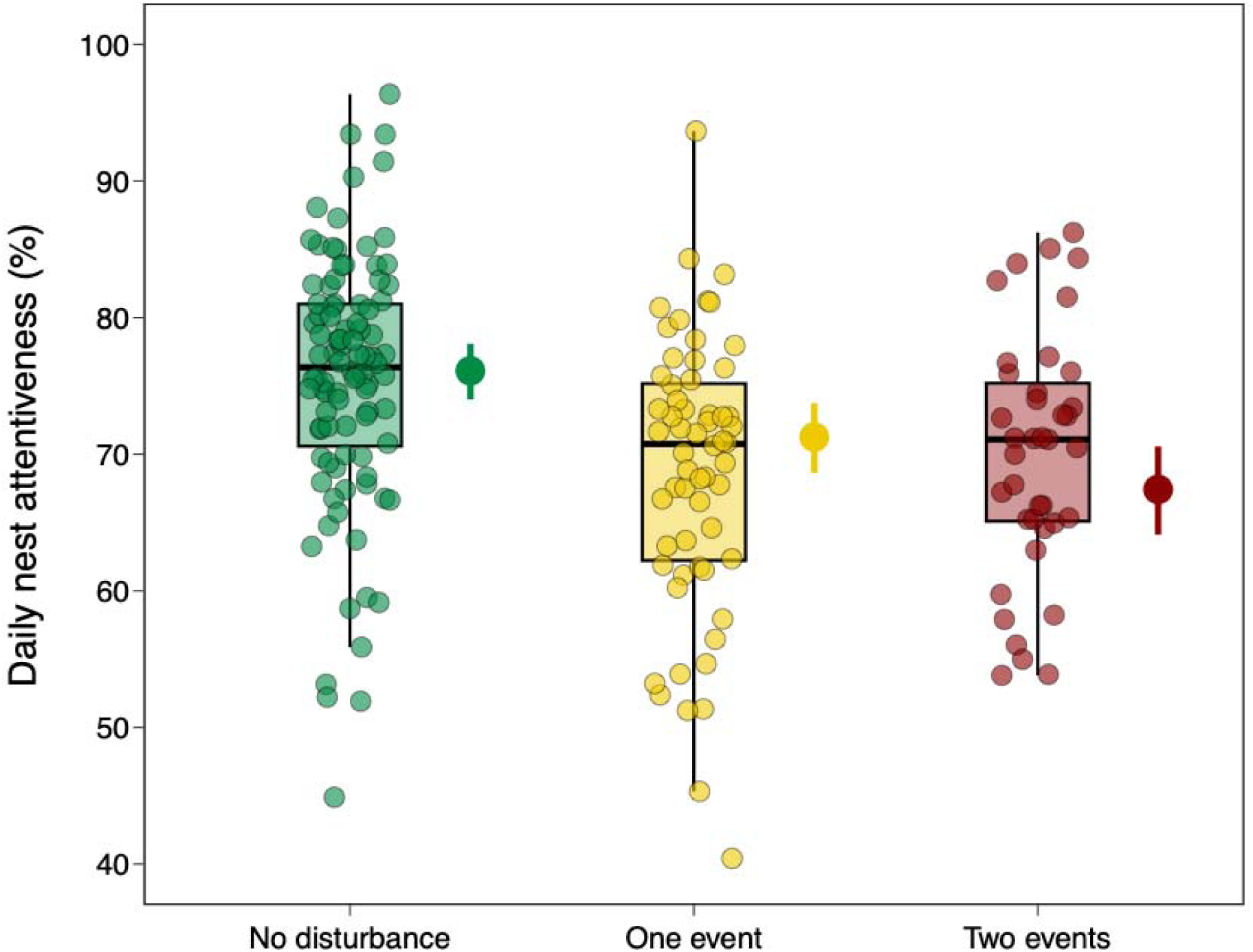
Daily nest attentiveness (proportion of time a female spends incubating compared to the time spent out of the nest on self-care related activities) depending on the number of disturbance events in a day. Box-plots represent the median and 50% of the values, and upper and lower whiskers represent values > 75th and < 25th percentile. Semi-transparent circles represent the bout duration raw data. Full circles and associated bars show the model estimates and 95% confidence intervals

**Table 3.** Estimates of a generalized linear mixed-effect model analysing the effect of the number of disturbance events on the daily nest attentiveness. Incubation day, daily ambient temperature, time of the day, population and year were also considered. Nest-box is a random effect. Significant effects P < 0.05 are highlighted in bold

|  | Estimate | SE | z | P |
| --- | --- | --- | --- | --- |
| Daily nest attentiveness |  |  |  |  |
| (R<sup>2</sup><sub>fixed factors</sub> = 0.33, R<sup>2</sup><sub>model</sub> = 0.68) |  |  |  |  |
| Intercept | 1.03 | 0.234 | 4.40 |  |
| **One event** | **-0.25** | **0.047** | **-5.36** | **<0.001** |
| **Two events** | **-0.43** | **0.063** | **-6.81** | **<0.001** |
| **Incubation day** | **0.19** | **0.047** | **3.97** | **<0.001** |
| **Temperature** | **-0.91** | **0.340** | **-2.68** | **0.007** |
| Temperature<sup>2</sup> | -0.31 | 0.318 | -0.98 | 0.327 |
| Time | -0.17 | 0.151 | -1.10 | 0.273 |
| Time<sup>2</sup> | -0.02 | 0.316 | -0.05 | 0.961 |
| Date | 0.15 | 0.129 | 1.20 | 0.229 |
| Population: Sagunto | 0.15 | 0.221 | 0.66 | 0.513 |
| **Year 2016** | **1.03** | **0.234** | **4.40** | **<0.001** |
| **Year 2017** | **-0.25** | **0.047** | **-5.36** | **<0.001** |

## Discussion

In this study, we found evidence that research disturbance alters Great tit incubation behaviour, keeping incubating females away from their nest. Our initial hypothesis was that females would react to a disturbance event by staying away from the nest longer than expected given their incubation rhythms, but we also expected that this anti-predatory behaviour, observed in previous studies (Kovařík and Pavel 2011; Ibáñez-Álamo and Soler 2012; Croston et al. 2021; Tkaczyk et al. 2023), was compensated with longer on-bouts that ended up balancing any negative effect on nest attentiveness. However, we found that 1) the response of females was restricted to the ongoing (absent females) or first (present females) off-bout after a disturbance, i.e., a short recovery period; 2) females did not clearly compensate for their prolonged absence from the nest by lengthening subsequent on-bouts; and consequently, and despite the limited anti-predator response, 3) daily incubation rhythms were negatively affected by the disturbance events (i.e., lower nest attentiveness).

Breeding females are flexible in their response to different perceived predation risks (Lima 2009; Martin and Briskie 2009). We found that females that were present in the nest at the time of a disturbance stayed away from the nest longer than absent females, presumably because they perceived being flushed as a greater risk than simply recognising an intruder from a distance. Flushed females also stayed away from the nest longer if the researcher performed additional activities during a visit (e.g., measuring eggs), in contrast to absent females, which seemed unaffected by prolonged visits. Experiments simulating the presence of predators near the nest generally result in incubating females leaving the nest for longer periods (Kovařík and Pavel 2011; Ibáñez-Álamo and Soler 2012; Croston et al. 2021; Tkaczyk et al. 2023), although the duration and magnitude of the response is species-specific (Ghalambor and Martin 2002; Zanette et al. 2011; Ghalambor et al. 2013) and/or dependent on predator type (Basso and Richner 2015a). Absent females appeared to evaluate the disturbance caused by the researchers differently. While flushed females remained near the nest-box, called and displayed mobbing behaviour before flying away (DDM pers. obs.), absent females may encounter a researcher in their nest when they return to incubate or while already foraging nearby. Flushed females may refrain from foraging or other self-care activities during part of their off-bout to increase vigilance (Morosinotto et al. 2013; Croston et al. 2021), at least until the researcher has left the nest, which could partly explain their longer off-bouts. An unexpected trend also emerged between populations when females were absent from the nest, although more data would be needed to assess a clear effect. In Sagunto, off-bouts of absent females tended to be shorter than in Pina. Given the anthropized nature of Sagunto, an extensive orange tree plantation, Great tit females could either be partially habituated to humans near the tree in which they built the nest, or the structure of the trees (short trees with crowns that generally reach the ground) would partly conceal the researcher from the foraging the bird.

Females were still constrained by a clutch that requires a suitable thermal range for embryo development (Drent 1975; DuRant et al. 2013). In elicited long absences from the nest, after being flushed, higher air temperature allowed females to stay off the nest for longer, pointing towards thermal constraints in the magnitude of the anti-predatory response, a trade-off between their own risk of being predated and the embryo survival that higher air temperature ameliorates. This is in line with a milder response when females were absent, where no association between air temperature and off-bout duration was found. Additionally, the general relationship between day of incubation and off-bout duration was expected (Diez-Méndez et al. 2021a) because when embryos develop their circulatory system, eggs cooling rates increase, causing a shortening in off-bouts (Cooper and Voss 2013).

Reduced activity near the nest, either by prolonged or less frequent off-bouts is the expected default behaviour when predation risk increases (Skutch 1957; Conway and Martin 2000). However, longer absences are usually compensated by longer incubation bouts, levelling out overall nest attendance (Kovařík and Pavel 2011; Ibáñez-Álamo and Soler 2012; Basso and Richner 2015b; Tkaczyk et al. 2023). In our study, despite the limited females’ response to a disturbance event, their daily incubation rhythms were altered, reducing their time on the nest. Reduced nest attendance may lengthen incubation period (Nord and Nilsson 2012) with consequences on egg hatchability (Macdonald et al. 2013; Diez-Méndez et al. 2020) in the short term, and reduce offspring survival in the long-term (Berntsen and Bech 2016; Ospina et al. 2018). Furthermore, Great tit females did not show a habituation response (see also Croston et al. 2021), and a second disturbance event within a day maintained the decline in daily nest attentiveness. Based on our results, it can be assumed that field experiments requiring regular, repeated visits during the incubation period may result in at least a decrease in egg hatchability compared to completely undisturbed nests. However, we were unable to link offspring productivity and egg hatchability to disturbance in our population, as all the nests were involved in different experiments and each of them was visited at least three days during the incubation period.

Our models showed a certain amount of variance that was not explained by the variables we measured. Physiological conditions at the individual level, which were not considered in this study, may explain some of the observed variability. For example, prolactin concentration following a disturbance may regulate nest attentiveness (Hope et al. 2020), and poorer body condition could lead to higher nest desertion rates (Dubiec 2011). Female personality could also explain the observed variability, as shy individuals take longer to return to the nest after being exposed to a novel object (Cole and Quinn 2014), although a long-term study on the effects of human recreation on Great tit breeding parameters showed no measurable within-population differences given their risk-taking personality (Hutfluss and Dingemanse 2019). Overall, Great tits are a resilient species that tolerates high degree of manipulation in the nest compared to other passerines (e.g., shrikes Tryjanowski and Kuźniak 1999; Golawski and Zduniak 2022). However, we must not lose sight of the importance of reducing disturbance from nest building to egg hatching. Visiting a nest while the female is absent is desirable, although less likely if it is not intentional, because incubating Great tit females sit on the nest for about 75% of their active day (Bueno-Enciso et al. 2017; Schöll et al. 2019; Diez-Méndez et al. 2021a). Handling females in the nest during incubation can lead to nest abandonment (Dubiec 2011) but nest visits also generate disturbances that disrupt incubation rhythms, a problem that should be taken into account when planning experimental approaches in the breeding populations.

## Conflicts of interest

The authors declare no competing interests.

## Ethical approval

This project was undertaken with the necessary ringing licences (no. 440135) and permits expedited by the regional (Generalitat Valenciana, Conselleria d’agricultura, medi ambient, canvi climàtic I desenvolupament rural) and the Spanish government (Ministry of Agriculture, Food and Environment). All procedures described in this project have complied with the guidelines of the University of Valencia.

## Data availability

Raw data and code for reproducibility are available in Zenodo (doi.org/10.5281/zenodo.10984081).

## Acknowledgments

We would like to thank all the collaborators and students for their assistance during the breeding seasons 2015-2017 in Sagunto and Pina, and specially José Verdejo for his help in Pina.

## Funding

This study was supported by projects CGL2013-48001-C2-1-P, CGL2016-79568-C3-1-P (former Spanish Ministry of Economy and Competitiveness, MINECO) and PID2021-122171NB-I00/ AEI/10.13039/501100011033/ FEDER, EU (Spanish Ministry of Science and Innovation). LC benefitted from a BRMIE (French regional grant for international student mobility), DD-M benefitted from an FPI fellowship (BES-2014-069191, MINECO) at the time, and is currently supported by the ERC (European Research Council) grant BABE 805189 and a GAČR Star grant (22-17593M).

## Notes

https://www.doi.org/10.5281/zenodo.10984081

### Competing Interest Statement

The authors have declared no competing interest.

### Summary of Updates

Introduction has been updated to clarify the background of the project and our hypotheses: Material and Methods has been updated for a better understanding of data collection and analysis; Results have been restructured for readability.

