## Supplementary material for "Research disturbance negatively impacts incubation behaviour of female Great tits"

**This is a pre-print of an article published in Behavioral Ecology and Sociobiology. The final authenticated version is available online at https://doi.org/10.1007/s00265-024-03514-y**

**Supplementary material**

**Research disturbance negatively impacts incubation behaviour of female Great tits**

**Léanne Clemencin^1^, Emilio Barba^2^ & David Diez-Méndez^3,4^***

^1^ Université Claude Bernard Lyon 1, 69622 Villeurbanne Cedex, France

^2^ “Cavanilles” Institute of Biodiversity and Evolutionary Biology, University of Valencia, Valencia, Spain.

^3^ Biology Centre of the Czech Academy of Sciences, Institute of Entomology, České Budějovice, Czech Republic.

^4^ University of South Bohemia, Faculty of Science, České Budějovice, Czech Republic.

* Corresponding author

**Table S1:** Descriptive statistics of off- and on-bouts length occurring around a disturbance event. It was taken into account whether a female was present in the nest or absent during the event.

|  | **Category** | **Median** | **Mean ± sd** | **Range** | **Sample size** |
| --- | --- | --- | --- | --- | --- |
| **Off-bouts (female absent)** | |  |  |  |  |
|  | Before | 10.2 | 15.1 ± 19.90 | 1.8 – 116 | 88 |
|  | During | 13.8 | 16.3 ± 11.20 | 3 – 93 | 88 |
|  | After | 9.9 | 11.8 ± 8.34 | 1.8 – 42.6 | 88 |
| **Off-bouts (female present)** | |  |  |  |  |
|  | Before | 9.6 | 11.6 ± 8.20 | 1.8 – 52.8 | 185 |
|  | After | 16.8 | 23.9 ± 20.90 | 3 – 119 | 185 |
|  | Next | 10.8 | 13.2 ± 10.60 | 1.8 – 79.8 | 185 |
| **On-bouts (female absent)** | |  |  |  |  |
|  | Before | 29.4 | 32.2 ± 18.80 | 6 – 90 | 90 |
|  | After | 30 | 37.1 ± 22.80 | 1.2 – 113 | 90 |
|  | Next | 27.3 | 34.5 ± 22.30 | 1.8 – 110 | 90 |
| **On-bouts (female present)** | |  |  |  |  |
|  | Before | 29.1 | 33.7 ± 22.00 | 4.2 – 139 | 178 |
|  | After | 30.9 | 35.8 ± 19.40 | 4.8 – 103 | 178 |
|  | Next | 31.5 | 37.1 ± 24.50 | 3 – 226 | 178 |

**Table S2:** Estimates of linear mixed-effect models analysing the effect of a disturbance event on the on-bout after a disturbance (female either absent or present in the nest-box) compared to the previous and next on-bouts. Incubation day, ambient temperature, time of the day, population and year were also considered. Nest-box is a random effect. Significant effects P < 0.05 are highlighted in bold.

|  |  | **Estimate** | **se** | ***t*** | ***P*** |
| --- | --- | --- | --- | --- | --- |
| On-bout (female absent) | |  |  |  |  |
| (R^2^ _fixed factors_ = 0.13, R^2^ _model_ = 0.42) | |  |  |  |  |
|  | Intercept | 3.40 | 0.314 | 10.83 |  |
|  | Previous | -0.11 | 0.086 | -1.25 | 0.213 |
|  | Next | -0.09 | 0.083 | -1.11 | 0.268 |
|  | Incubation day | 0.11 | 0.073 | 1.52 | 0.134 |
|  | Temperature | 1.15 | 0.782 | 1.47 | 0.144 |
|  | Temperature^2^ | 0.18 | 0.685 | 0.27 | 0.790 |
|  | Time | -1.01 | 0.760 | -1.32 | 0.187 |
|  | Time^2^ | -0.30 | 0.823 | -0.37 | 0.714 |
|  | Date | -0.27 | 0.190 | -1.42 | 0.166 |
|  | Population: Sagunto | -0.61 | 0.354 | -1.72 | 0.096 |
|  | **Year 2016** | **0.65** | **0.242** | **2.70** | **0.011** |
|  | Year 2017 | 0.43 | 0.238 | 1.80 | 0.082 |
| On-bout (female present) | |  |  |  |  |
| (R^2^ _fixed factors_ = 0.04, R^2^ _model_ = 0.23) | |  |  |  |  |
|  | Intercept | 3.26 | 0.268 | 12.16 |  |
|  | Previous | -0.12 | 0.061 | -1.92 | 0.055 |
|  | Next | 0.01 | 0.057 | 0.17 | 0.865 |
|  | Incubation day | 0.03 | 0.047 | 0.62 | 0.537 |
|  | Temperature | 0.75 | 0.763 | 0.98 | 0.327 |
|  | Temperature^2^ | -0.12 | 0.614 | -0.19 | 0.849 |
|  | Time | -1.22 | 0.724 | -1.68 | 0.094 |
|  | Time^2^ | 0.12 | 0.686 | 0.18 | 0.860 |
|  | Date | 0.01 | 0.118 | 0.11 | 0.912 |
|  | Population: Sagunto | 0.24 | 0.194 | 1.24 | 0.223 |
|  | Year 2016 | 0.06 | 0.187 | 0.34 | 0.735 |
|  | Year 2017 | 0.08 | 0.230 | 0.25 | 0.802 |

**Table S3:** Estimates of a linear mixed-effect model analysing the effect of the duration of a disturbance event (short vs. long disturbance) on the duration of the ongoing off-bout when a female was absent the nest-box. Incubation day, ambient temperature, time of the day, population and year were also considered. Nest-box is a random effect.

|  |  | **Estimate** | **se** | ***t*** | ***P*** |
| --- | --- | --- | --- | --- | --- |
| Off-bout duration (female absent) | |  |  |  |  |
| (R^2^ _fixed factors_ = 0.14, R^2^ _model_ = 0.17) | |  |  |  |  |
|  | Intercept | 2.99 | 0.267 | 1.12 |  |
|  | Long disturbance | 0.12 | 0.184 | 0.69 | 0.492 |
|  | Incubation day | 0.00 | 0.082 | 0.01 | 0.989 |
|  | Temperature | -0.21 | 0.635 | -0.33 | 0.739 |
|  | Temperature^2^ | 0.21 | 0.622 | 0.34 | 0.736 |
|  | Time | -0.31 | 0.616 | -0.50 | 0.619 |
|  | Time^2^ | -0.67 | 0.708 | -0.95 | 0.346 |
|  | Date | -0.12 | 0.163 | -0.72 | 0.477 |
|  | Population: Sagunto | -0.58 | 0.303 | -1.93 | 0.060 |
|  | Year 2016 | 0.12 | 0.228 | 0.52 | 0.606 |
|  | Year 2017 | -0.00 | 0.226 | -0.02 | 0.986 |
